# Guidelines for reliable and reproducible functional enrichment analysis

**DOI:** 10.1101/2021.09.06.459114

**Authors:** Kaumadi Wijesooriya, Sameer A Jadaan, Kaushalya L Perera, Tanuveer Kaur, Mark Ziemann

## Abstract

Gene set enrichment tests (a.k.a. functional enrichment analysis) are among the most frequently used methods in computational biology. Despite this popularity, there are concerns that these methods are being applied incorrectly and the results of some peer-reviewed publications are unreliable. These problems include the use of inappropriate background gene lists, lack of false discovery rate correction and lack of methodological detail. An example analysis of public RNA-seq reveals that these methodological errors alter enrichment results dramatically. To ascertain the frequency of these errors in the literature, we performed a screen of 186 open access research articles describing functional enrichment results. We find that 95% of analyses using over-representation tests did not implement an appropriate background gene list or did not describe this in the methods. Failure to perform p-value correction for multiple tests was identified in 43% of analyses. Many studies lacked detail in the methods section about the tools and gene sets used. Only 15% of studies avoided major flaws, which highlights the poor state of functional enrichment rigour and reporting in the contemporary literature. We provide a set of minimum standards that should act as a checklist for researchers and peer-reviewers.

## Background

Since the turn of the millennium, high throughput “omics” techniques like microarrays and high throughput sequencing have brought with them a deluge of data. These experiments involve the measurement of thousands of genes simultaneously, and can identify hundreds or even thousands of significant associations with developmental stages or diseases. Interpreting such data is extraordinarily challenging, as the sheer number of associations can be difficult to investigate in a gene-by-gene manner. Instead, many tools have been developed in an effort to summarise gene profiles into simplified functional categories. These functional categories typically represent signaling or biochemical pathways, curated from information present in the literature, hence the name functional enrichment.

Widely used functional enrichment tools can be classified into two main categories; (i) over-representation analysis (ORA) and (ii) functional class scoring (FCS), and the most common application is in differential gene expression analysis. In ORA, differentially expressed genes (DEGs) meeting a significance and/or fold change threshold are queried against curated pathways (gene sets). A statistical test is performed to ascertain whether the number of DEGs belonging to a particular gene set is higher than that expected by random chance, as determined by comparison to a background gene list. These ORA tools can be stand-alone software packages or web services, and they use one or more statistical tests (eg: Fisher’s Exact test, hypergeometric test) **[1,2]**.

In the case of ORA for differential expression (eg: RNA-seq), a whole genome background is inappropriate because in any tissue, most genes are not expressed and therefore have no chance of being classified as DEGs. A good rule of thumb is to use a background gene list consisting of genes detected in the assay at a level where they have a chance of being classified as DEG **[3]**.

FCS tools involve giving each detected gene a differential expression score and then evaluating whether the scores are more positive or negative than expected by chance for each gene set. The popular Gene Set Enrichment Analysis (GSEA) tool uses permutation approaches to establish whether a gene set is significantly associated with higher or lower scores, either by permuting sample labels or by permuting genes in the differential expression profile **[4]**.

From a user’s perspective, ORA is easier to conduct because it is as simple as pasting a list of gene names into a text box on a website. FCS tools are more difficult to use, but they are more sensitive at detecting subtle associations **[5–7]**.

Although these are powerful tools to summarise complex genomics data, there are concerns that they are not being correctly used. Tipney and Hunter **[3]** warn that inappropriate background set selection heavily influences enrichment results. Timmons et al **[8]** highlight two cases where an inappropriate background list led to invalid enrichment results in published articles.

The purpose of this work is to survey the frequency of methodological and reporting flaws in the literature, in particular; (i) inappropriate background gene set, (ii) lack of p-value adjustment for multiple comparisons and (iii) lack of essential methodological details. We use the results of this survey to inform a set of minimum standards for functional enrichment analysis.

## Results

### An example of functional enrichment analysis misuse

To demonstrate the effect of functional enrichment analysis misuse, we used an example RNA-seq dataset examining the effect of high glucose exposure on hepatocytes. Out of 39,297 genes in the whole genome, 15,635 were above the detection threshold (≥10 reads per sample on average). Statistical analysis revealed 3,472 differentially expressed genes with 1,560 up-regulated and 1,912 down-regulated (FDR<0.05) due to elevated glucose exposure.

FCS revealed 95 up and 316 down regulated gene sets (FDR<0.05). ORA with a background gene list consisting of detected genes revealed 55 upregulated and 238 downregulated gene sets (FDR<0.05). The overlap between FCS and ORA shown in Fig. 1a is reflected by a Jaccard statistic of 0.66, indicating moderate concordance between these methods.

**Figure 1.**
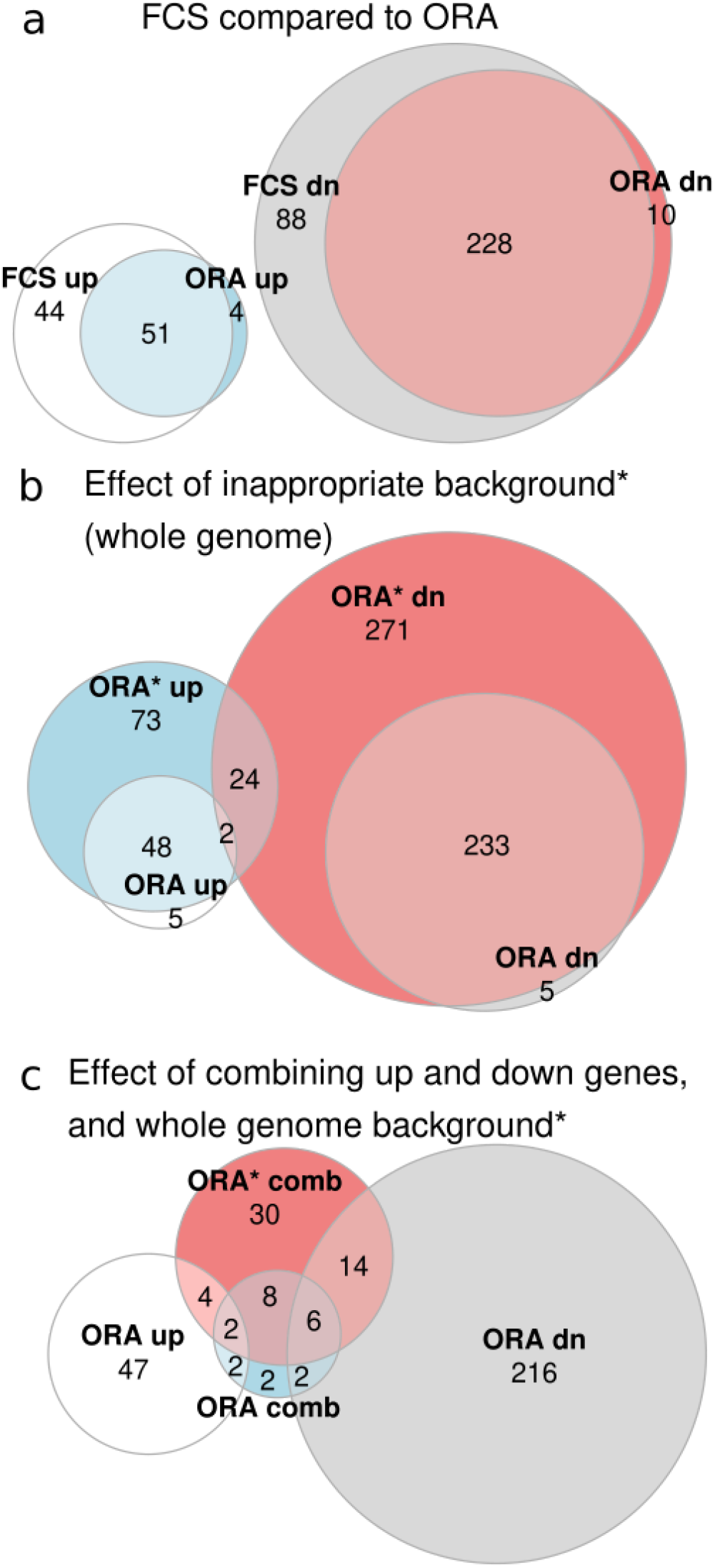
Example of enrichment analysis misuse. Euler diagrams show the overlap between sets of differentially regulated gene sets with different methods (FDR<0.05). **a** Comparison of FCS and ORA methods. **b** ORA with standard and whole genome background. Whole genome background analysis is indicated with a *. **c** Effect of combining up and downregulated genes on ORA results. Analysis using combined gene list is indicated by ORA comb, while ORA* comb indicates combined gene list analysis with a whole genome background.

We then performed ORA using the whole genome gene list (indicated as ORA*), which resulted in 147 up and 530 down regulated gene sets (FDR<0.05), The overlap of ORA with ORA* was relatively small, with a Jaccard statistic of 0.41 (Fig. 1b). Interestingly, 26 gene sets were simultaneously up and downregulated with this approach.

Then we performed ORA analysis after combining up and downregulated genes as sometimes observed in the literature (indicated as ORA comb) with the standard background and again with a whole genome background (indicated as ORA* comb). ORA comb revealed only 22 differentially regulated gene sets with a Jaccard similarity to ORA of only 0.04 (Fig. 1c). ORA* comb yielded 64 differentially regulated gene sets with a Jaccard similarity to ORA of only 0.08. These results suggest that when used properly, FCS and ORA yield similar results, but when used improperly, ORA results can be severely distorted.

### A screen of functional enrichment analyses in open access journals

A search of PubMed Central showed 2,941 articles published in 2019 with the keywords “enrichment analysis”, “pathway analysis” or “ontology analysis”. From these, we initially selected 200 articles for detailed methodological analysis. We excluded 14 articles from the screen because they did not present any enrichment analysis. Those excluded articles included articles describing novel enrichment analysis techniques or tools, review articles or conference abstracts. As some articles included more than one enrichment analysis, the dataset included 235 analyses from 186 articles; this data is available in Table S1. A flow diagram of the survey is provided in Fig. 2.

**Figure 2.**
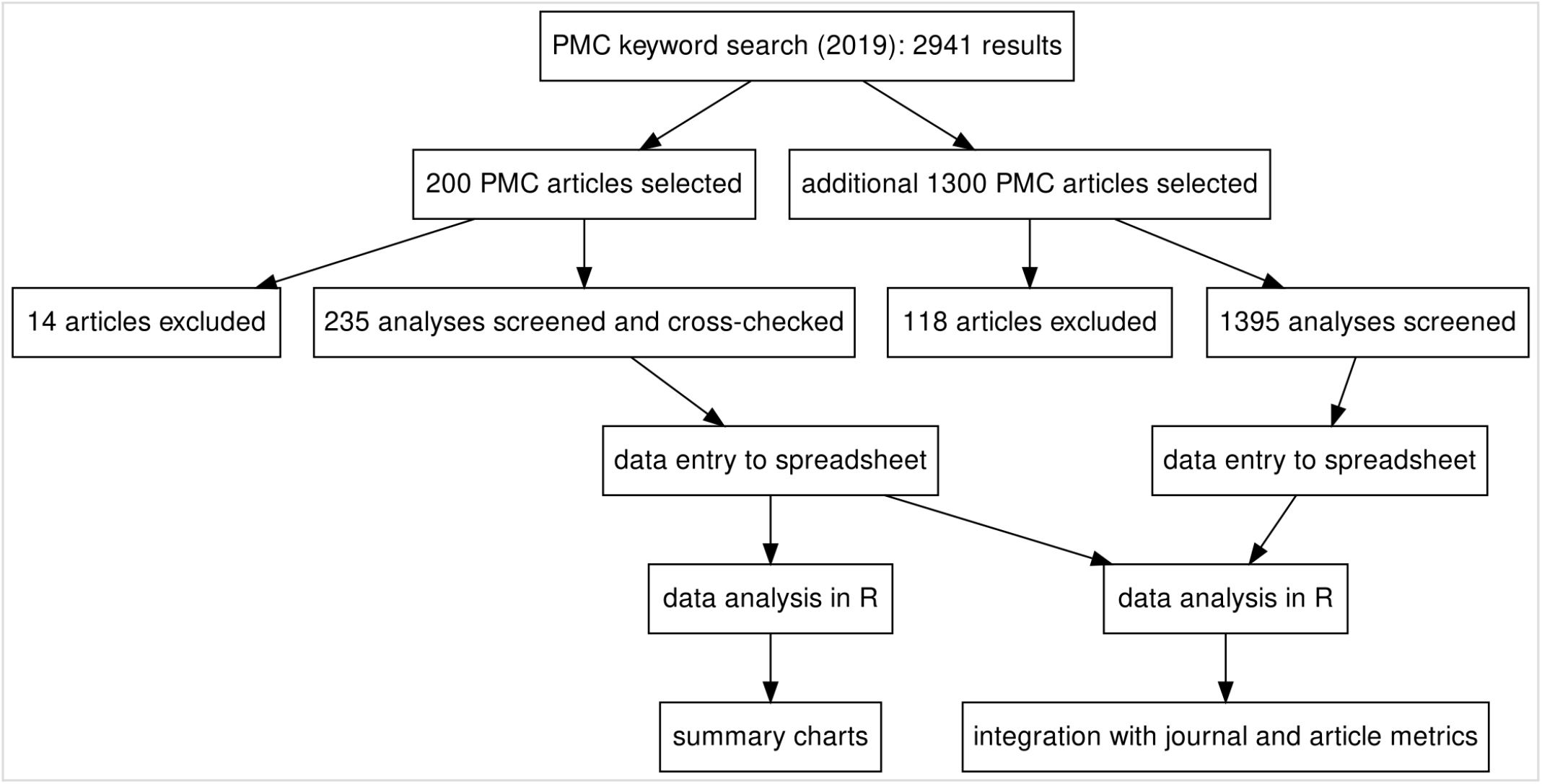
A summary of the survey of functional enrichment analyses. The survey consists of two parts, with 200 and 1300 PMC articles considered respectively.

There were articles from 96 journals in the sample, with *PLoS One*, *Scientific Reports*, and *PeerJ* being the biggest contributors (Figure S1a). There were 18 different omics types, with gene expression array and RNA-seq being the most popular (Figure S1b). There were 31 different species under study, but *Homo sapiens* was the most common with 157 analyses (Figure S1c).

We recorded the use of 26 different gene set libraries, with GO and KEGG being the most frequently used (Fig. 3a). There were 14 analyses where the gene set libraries used were not defined in the article. Only 18 analyses reported the version of the gene set library used (Fig. 3b). There were 12 different statistical tests used, and the most common reported tests were Fisher’s Exact, GSEA and hypergeometric tests; but the statistical test used was not reported for the majority of analyses (Fig. 3c). Fourteen analyses did not conduct any statistical test, they only showed the number of genes belonging to different sets. Out of the 225 analyses that performed a statistical test, only 119 (53%) described correction of p-values for multiple testing (Fig 3d).

**Figure 3.**
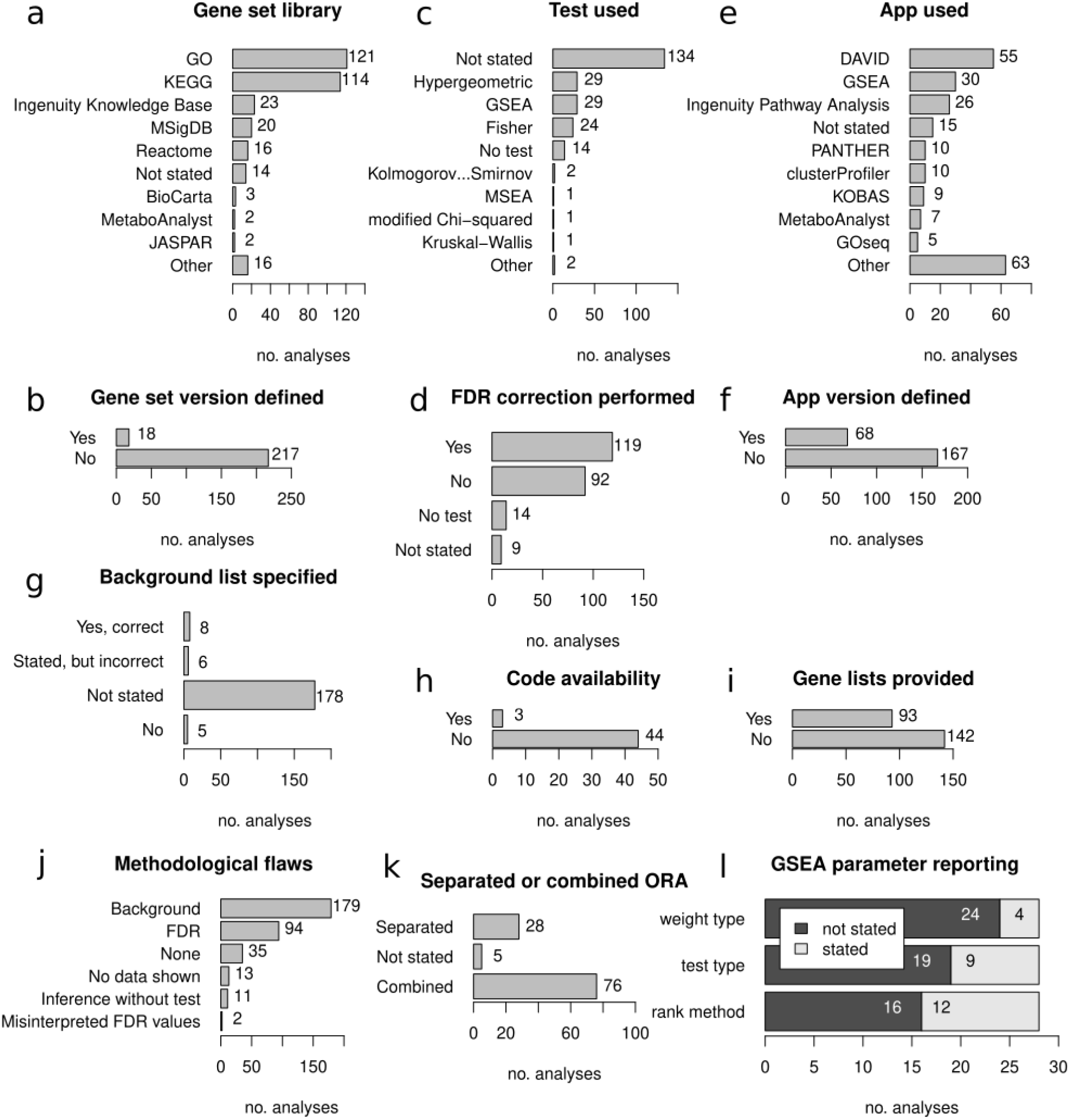
Findings from the survey of published enrichment analyses. **a** Representation of different gene set libraries. **b** Proportion of analyses reporting gene set library version information. **c** Representation of different statistical tests. **d** Proportion of analyses conducting correction of p-values for multiple comparisons. **e** Representation of different software tools for enrichment analysis. **f** Proportion of analyses that reported version information for software used. **g** Background gene set usage and reporting. **h** Proportion of scripted analyses that provided software code. **i** Proportion of analyses that provided gene profiles. **j** Proportion of analyses with different methodological problems. **k** Whether up- and down-regulated genes were analysed separately or combined. Limited to ORA of differential expression only. **l** Reporting of GSEA parameters.

There were 50 different tools used to perform enrichment analysis, with DAVID and GSEA being the most common; while 15 analyses (6.4%) did not state what tool was used (Fig. 3e). The version of the software used was provided in only 68 of 235 analyses (29%) (Fig. 3f).

For analyses using ORA methods, we examined what background gene set was used (Fig. 3g). This revealed that in most cases, the background list was not defined, or it was clear from the article methods section that no background list was used. In a few cases, a background list was mentioned but was inappropriate, for example using a whole genome background for an assay like RNA-seq. In only 8/197 of cases (4.1%), the appropriate background list was described in the article.

Of the 47 analyses which used a scripted computer language, only 3 provided links to the code used for enrichment analysis (6.4%) (Fig. 3h). For 93 of 235 analyses (40%), the corresponding gene lists/profiles were provided either in the supplement or in the article itself (Fig. 3i).

Next, we quantified the frequency of methodological issues and reporting that would undermine the conclusions (Fig. 3j). Lack of appropriate background was the most common issue (179 cases), followed by FDR (94 cases), then lack of data shown (13), inference without test (11 cases), and misinterpreted FDR values (2 cases). Only 35 analyses (15%) did not exhibit any of these major methodological issues.

During this survey, we noticed some studies grouped up- and down-regulated gene lists together prior to ORA, a practice we were not expecting. To assess this in a systematic way we assessed differential gene expression studies to determine how common it was to combine up- and down-regulated gene lists prior to ORA (Fig 3k). From 109 analyses, just 28 studies performed separate analysis of up- and down-regulated gene sets.

We also looked at studies performing GSEA, and whether three important analytical choices were described in the methods. These are (i) the gene weighting parameter, (ii) test type, ie: permuting sample labels or genes, and (iii) method used for ranking genes. These parameters were not stated in more than half of the analyses (Fig. 3l).

Taken together, these data suggest methodological and reporting deficiencies are widespread in published functional enrichment analyses.

### Analysis scores

To make an accurate assessment of association between analysis quality and bibliometrics, we required a larger sample of articles. We therefore analysed a further 1,300 articles, bringing the total number of analyses described up to 1630; this dataset is available in Table S2.

We then scored each analysis based on the presence or absence of methodological issues and included details. The median score was −4, with a mean of −3.5 and standard deviation of 1.4 (Fig. 4a). Next, we assessed whether these analysis scores are associated with Scimago Journal Rank (SJR), a journal-level citation metric (Fig. 4b). There was no statistically significant association of analysis score and SJR (Pearson r=0.03, p=0.23).

**Figure 4.**
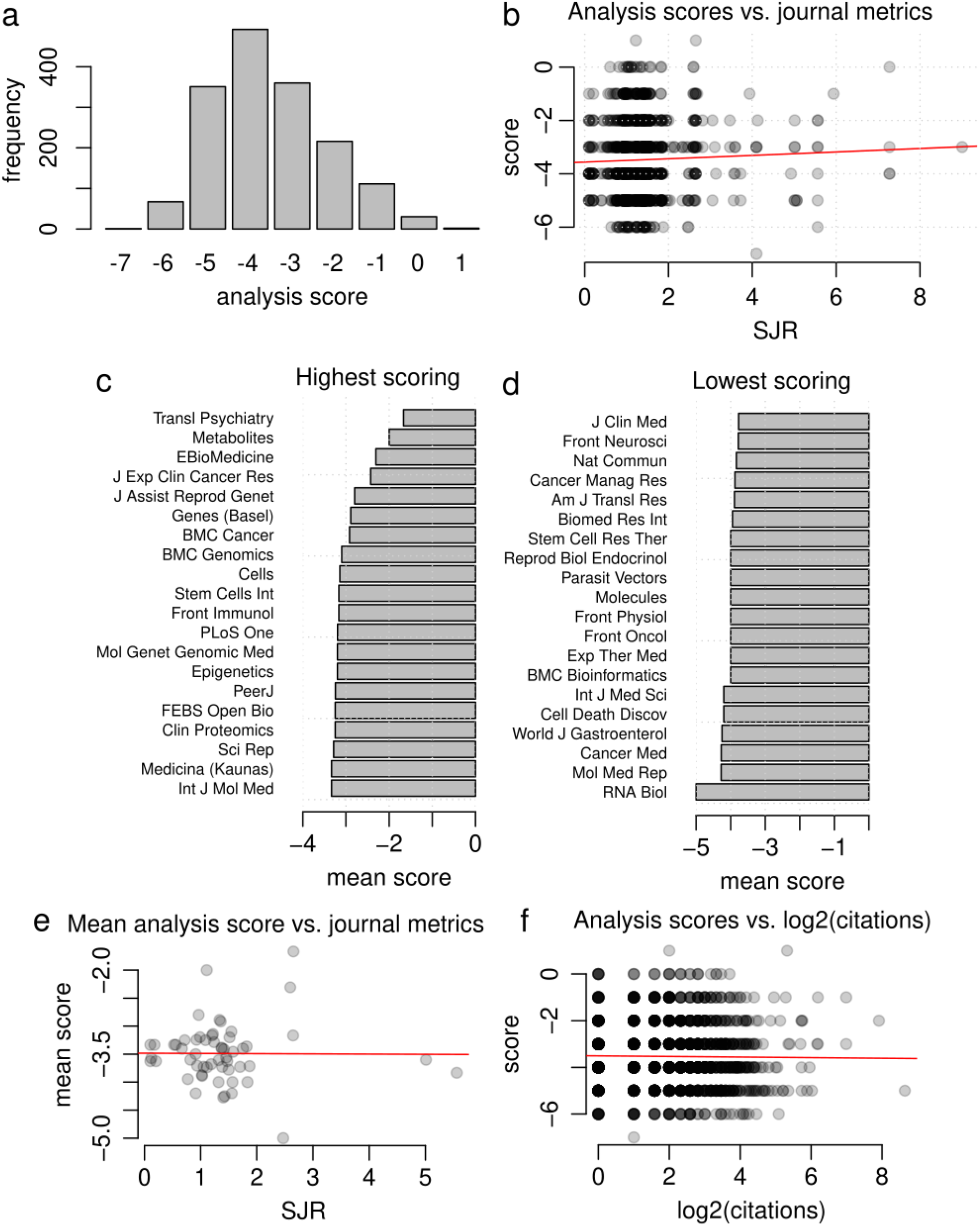
Comparison of analysis scores to bibliometrics. **a** Distribution of analysis scores. **b** Association of analysis scores with Scimago Journal Rank (SJR). **c** Journals with highest mean analysis scores. **d** Journals with lowest mean analysis scores. **e** Association of mean analysis score with SJR for journals with 5 or more analyses. **f** Association of analysis scores with accrued citations.

Next, we wanted to know which journals had the highest and lowest scores. Only journals with five or more analyses were included. The best scoring journals were *Transl Psychiatry*, *Metabolites* and *J Exp Clin Cancer Res* (Fig. 4c), while the poorest were *RNA Biol*, *Mol Med Rep*, *Cancer Med* and *World J Gastroenterol* (Fig. 4d), although we note that there was a wide variation between articles of the same journal.

Then we assessed for an association between mean analysis score and the SJR, for journals with 5 or more analyses (Fig. 4e). Again, there was no association between analysis score and SJR (Pearson r=−0.008, p=0.95). Next, we assessed whether there was any association between analysis scores and the number of citations received by the articles. After log transforming the citation data, there was no association between citations and analysis scores (Pearson r=0.002, p=0.95) (Fig. 4f). Taken together, these findings suggest that these methodological issues are not limited to lower ranking or poorly cited articles.

## Discussion

While there have been several articles benchmarking the performance of functional enrichment tools, there has been relatively little written on how existing tools are used (or misused). A recent study looked at the popularity of frequently used tools and benchmarked their performance **[7]**, but this assumes that the tools were properly used, and methods sufficiently reported. In this sample of open access research articles, there appears to be a bias toward tools that are easy to use. ORA tools that only require pasting lists of gene identifiers into a webpage (ie: DAVID, KOBAS and PANTHER) are collectively more popular than other solutions like GSEA (a stand-alone graphical user interface software for FCS) or any command line tool. This is despite the fact that ORA tools perform worse than FCS and new generation pathway topology tools **[5–7]**.

Failing to properly describe the background gene list was the most common methodological issue (Fig. 3j). This error can lead to substantially distorted results (Fig. 1b). In the example here, using the inappropriate whole genome background resulted in 2.3-fold more differentially regulated gene sets. Thus, if there are few or no significant results using a correct background list, there may be a temptation to use an inappropriate background to avoid null results.

The proportion of ORA studies that combined up- and down-regulated gene lists prior to analysis was high, at 70% (Fig. 3k). As exemplified in Fig 1c, combining up- and down-regulated genes can severely reduce the number of significant gene sets (from 293 gene sets to just 22 in this example). This is consistent with a previous report indicating that combining up- and down-regulated gene lists is considered poor practice because it reduces sensitivity **[9]**. These results suggest that a large fraction of published works suffer from a high rate of false negatives, in effect missing a large fraction of truly differentially regulated gene sets. This is exacerbated by using an inappropriate background gene list, which leads to a high rate of false positives.

Although analyses involving GSEA scored better overall, they were not free of issues. For example, GSEA has different options that impact the results including the ranking metric, gene weighting method and the permutation type (on samples or genes), which were not regularly reported in articles (Fig. 3l), limiting reproducibility.

We scored a total of 1630 analyses, revealing only a small fraction of articles obtained a satisfactory score of zero or higher. The analysis scores we generated did not correlate with journal or article metrics. This suggests that methodological and reporting problems are not limited to lower ranked journals, but are a more general problem.

Still there are some limitations that need to be recognised. Many open access articles examined here are from lower-ranked journals that might not be representative of articles in paywalled journals. The articles included in this study contained keywords related to functional enrichment in the abstract, and it is plausible that articles in higher ranked journals contain such details in the abstract at lower rates.

Nevertheless, these results are a wake-up call for reproducibility and highlight the urgent need for minimum standards for functional enrichment analysis. To address this, we provide a set of minimum standards for functional enrichment analysis, as well as additional recommendations to achieve gold standard reliability and reproducibility (Box 1 and 2).

### Box 1.

**Essential minimum standards for functional enrichment analysis**

The following guidelines are selected to avoid the most severe methodological issues while requiring minimal effort to address.

1. Before starting the analysis, read the tool’s user manual.
2. Report the origin of the gene sets and the version used or date downloaded. These databases are regularly upgraded so this is important for reproducibility.
3. Report the tool used and its software version. As these tools are regularly updated, this will aid reproducibility. For online tools, record the date the tool was used.
4. If making claims about the regulation of a gene set, then results of a statistical test must be shown including measures of significance and effect size. The number of genes belonging to a pathway on its own is insufficient to infer enrichment.
5. Report which statistical test was used. This will help long term reproducibility, for example if in future the tool is no longer available. This is especially relevant for tools that report the results of different statistical tests.
6. Always report FDR or q-values (p-values that have been corrected for multiple tests). This will reduce the chance of false positives when performing many tests simultaneously [10]. Report what method was used for p-value adjustment. Bonferroni, FDR and FWER are examples.
7. If ORA is used, it is vital to use a background list consisting of all detected genes, and report this in the methods section. Avoid tools that don’t accept a background list.
8. Report selection criteria and parameters. If performing ORA, then the inclusion thresholds need to be mentioned. If using GSEA or another FCS tool, parameters around ranking, gene weighting and type of test need to be disclosed (eg: permuting sample labels or gene labels). Any parameters that vary from the defaults should be reported.
9. If using ORA for gene expression analysis, be sure to conduct ORA tests separately for up- and down-regulated genes before analysis. Combining up- and down-regulated genes into a single list will limit sensitivity.
10. Functional enrichment analysis results shouldn’t be considered proof of biological plausibility nor validity of omics data analysis [8]. Such tools are best used to generate new hypotheses; informing subsequent biochemical or signaling experimentation.

### Box 2.

**Additional recommendations for enhanced reliability and reproducibility**

The following guidelines are directed at going over and above the minimum standards towards a gold standard of reliability and reproducibility.

11. Preference FCS and pathway topology tools over ORA tools. ORA tools have a lower sensitivity. In practice, they are rarely used properly.
12. Include the gene profile data (including any background lists) in the supplementary data in TSV or CSV formats. Excel spreadsheets are not recommended for genomics work **[11]**.
13. Scripted analysis workflows are preferred over analysis involving graphical user interfaces because they can attain a higher degree of computational reproducibility, so consider investing in upgrading your analysis. Similarly, free and open-source software tools are preferred over proprietary software.
14. If using a scripted analysis, provide code by depositing it to a repository like GitHub or Zenodo. Link the data to the code, so anyone can download and reproduce your results **[12]**.

## Methods

### An example gene set analysis

To demonstrate the effect of misusing gene set analysis, a publicly available RNA-seq dataset (SRA accession SRP128998) was downloaded from DEE2 on 13th August 2021 **[13]**. This data consists of immortalised human hepatocytes cultured in standard (n=3) or high glucose (n=3), first described by Felisbino et al **[14]**. Transcript level counts were aggregated to genes using the getDEE2 R package v1.2.0. Next, genes with an average of less than 10 reads per sample were omitted from downstream analysis. Differential expression statistical analysis was conducted with DESeq2 v1.32.0 **[15]** to identify genes altered by high glucose exposure. For gene set analysis, human Reactome gene sets **[16]** were downloaded in GMT format from the Reactome website (accessed 13th August 2021). FCS was performed using the mitch R package v1.4.0 with default settings, which uses a rank-ANOVA statistical test **[5]**. Genes with FDR<0.05 were used for ORA analysis using the clusterProfiler R package (v4.0.2) enricher function that implements a hypergeometric test **[17]**. No fold-change threshold was used to select genes for ORA. For ORA, two types of background gene sets were used: (i) detected genes, or (ii) all genes in the genome. For genes and gene sets, a false discovery rate adjusted p-value (FDR) of 0.05 was considered significant. Analyses were conducted in R version 4.1.0.

### Survey of enrichment analysis

We collated 2,941 articles in PubMed Central published in 2019 that have keywords “enrichment analysis”, “pathway analysis” or “ontology analysis”. We initially sampled 200 of these articles and collected the following information from the article, searching the methods sections and other parts of the article including the supplement.

- Journal name
- Type of omics data
- Gene set library used, and whether a version was reported
- Statistical test used
- Whether p-values were corrected for multiple comparisons
- Software package used, and whether a version was reported
- Whether an appropriate background gene set was used
- Code availability
- Whether gene profile was provided in the supplement
- Whether the analysis had any major flaws that might invalidate the results. This includes:

i. background gene set not stated or inappropriate background set used,
ii. lack of FDR correction,
iii. no enrichment data shown,
iv. inference without performing any statistical test, and
v. misinterpreting p-values by stating results were significant when FDR values indicate they aren’t.

We excluded articles describing novel enrichment analysis techniques/tools, review articles and conference abstracts. Some articles presented the results of >1 enrichment analysis, so additional rows were added to accommodate them. These data were entered into a Google Spreadsheet by a team of five researchers. These articles were cross checked by another team member and any discrepancies were resolved.

For analyses using GSEA, we scanned the articles to identify whether key methodological steps were described, including (i) the gene weighting parameter, (ii) test type, ie: permuting sample labels or genes, and (iii) method used for ranking genes.

For ORA of differential gene expression analyses, we inspected the articles to see evidence that researchers were conducting enrichment of up and down regulated genes separately or combined.

Cleaned data were then loaded into R for downstream analysis.

For assessment of enrichment analysis quality with journal metrics and citations, we required a larger sample, so we selected a further 1300 articles from PMC for analysis. Results from this sample were not double-checked and so may contain a small number of inaccuracies. We rated each analysis with a simple approach that deducted points for missing methodological details and awarding points for including extra information (Table 1).

**Table 1.**
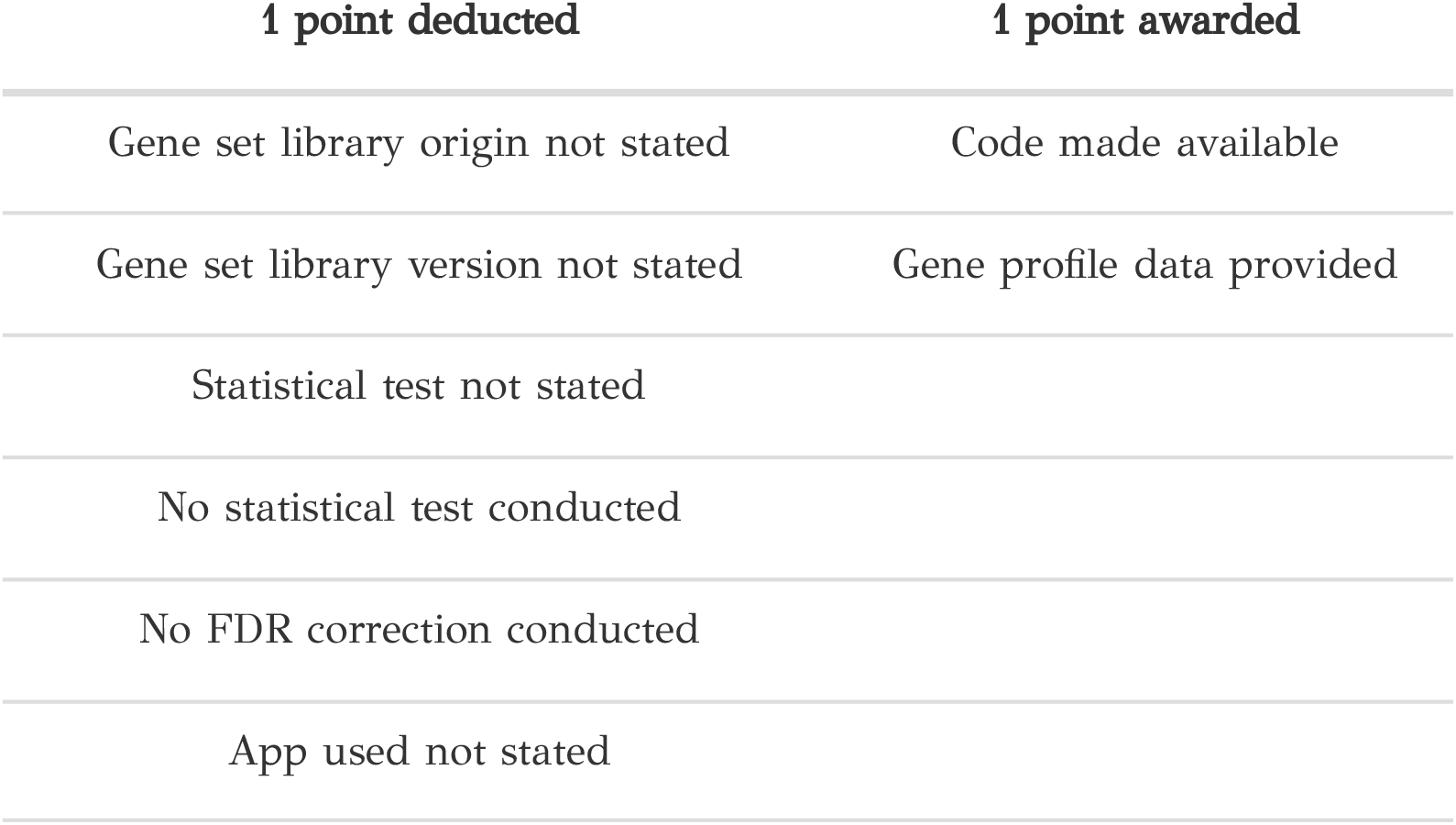
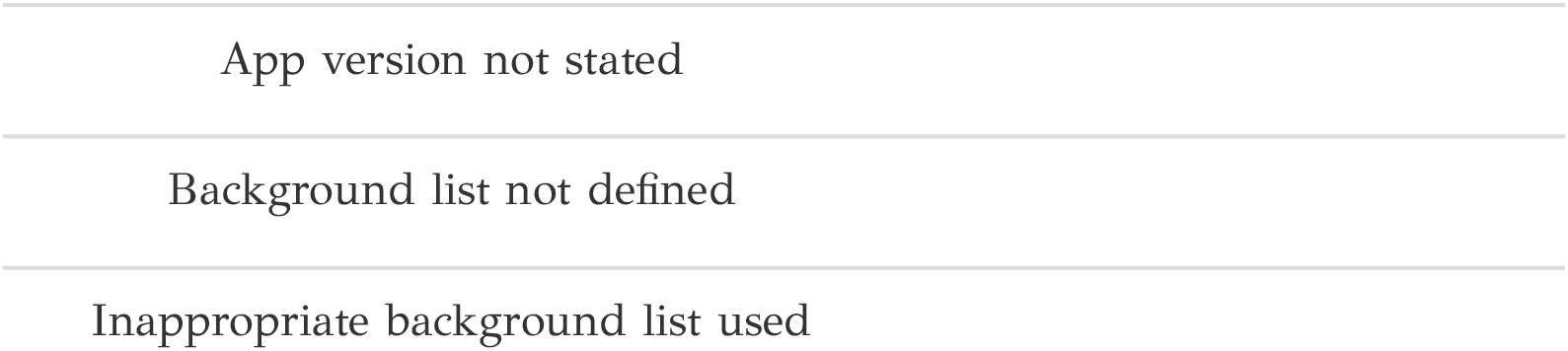
Scoring schema.

| **1 point deducted** | **1 point awarded** |
| --- | --- |
| Gene set library origin not stated | Code made available |
| Gene set library version not stated | Gene profile data provided |
| Statistical test not stated |  |
| No statistical test conducted |  |
| No FDR correction conducted |  |
| App used not stated |  |

SJR data for 2020 were downloaded from the Scimago website (accessed 5th August 2021) and used to score journals by their citation metrics. Using NCBI’s Eutils API, we collected the number of citations each article accrued since publication. In R, we used Pearson correlation tests to assess the association with the analysis scores we generated.

## Supplementary information

Additional file 1: Table S1. A survey of 186 articles describing functional enrichment results in TSV format.

Additional file 1: Table S2 A survey of 1365 articles describing functional enrichment results in TSV format.

## Declarations

### Ethics approval and consent to participate

Not applicable.

### Consent for publication

Not applicable.

### Availability of data and materials

The code and datasets described in this study are available from GitHub (https://github.com/markziemann/SurveyEnrichmentMethods).

### Authors’ contributions

Conceptualisation and design: MZ. Data acquisition and analysis: KW, SAJ, TK, KLP, MZ. Software: KW, MZ. Initial manuscript draft: KW, MZ. Manuscript editing: KW, SAJ, MZ. All authors read and approved the final manuscript.

## Acknowledgements

We thank Antony Kaspi (Walter and Eliza Hall Institute), Nick Wong and Anup Shah (Monash University) for comments on the manuscript. This research was supported by use of the Nectar Research Cloud, a collaborative Australian research platform supported by the NCRIS-funded Australian Research Data Commons (ARDC).

